# Conservation of gene expression patterns between the amniotic epithelium at birth and a newborn’s nasal epithelium

**DOI:** 10.1101/2024.08.22.609087

**Authors:** David G Hancock, Patricia Agudelo-Romero, Elizabeth Kicic-Starcevich, James Lim, Thomas Iosifidis, Yuliya V Karpievitch, Guicheng Zhang, David J Martino, Anthony Kicic, Stephen M Stick

**Affiliations:** Wal-yan Respiratory Research Centre, Telethon Kids Institute, University of Western Australia, Nedlands, 6009, Western Australia, Australia; Department of Respiratory and Sleep Medicine, Perth Children’s Hospital, Nedlands, 6009, Western Australia, Australia; School of Population Health, Curtin University, Bentley, 6102, Western Australia, Australia.; Centre for Cell Therapy and Regenerative Medicine, School of Medicine, The University of Western Australia, Nedlands, 6009, Western Australia, Australia; European Virus Bioinformatics Centre, Jena, Thuringia, Germany; School of Biomedical Sciences, The University of Western Australia, Nedlands, 6009, Western Australia, Australia

**Keywords:** amnion, nasal, respiratory, transcriptomics, network, surrogate

## Abstract

**Background:** Amniotic epithelial cells are foetal-derived stem cells, capable of differentiating into all three germ layers, including mature epithelial cell populations. However, the conservation between amniotic epithelium and other epithelial tissues has not been sufficiently explored. We hypothesised that the amniotic epithelium might serve as a surrogate tissue source for investigating transcriptional profiles in the respiratory epithelium of the newborn. We compared gene expression profiles and weighted gene co-expression network structure in paired amniotic and newborn nasal epithelial samples from 85 participants in the Airway Epithelium Respiratory Illnesses and Allergy (AERIAL) birth cohort.

**Results:** In total, 11,867 genes (79.7%) were commonly expressed in both amniotic and nasal epithelium, with uniquely expressed genes (2,563 and 458, respectively) enriched for biological functions related to each tissue’s specialist functions (e.g. developmental programs and ciliated cells, respectively). We observed a strong overlap in weighted gene co-expression network structure between both tissues, with ten co-expression modules identified in consensus network analysis. Genes commonly expressed in both tissues and/or found in the consensus network modules were enriched for biologically relevant gene signatures and pathway terms related to airway function. We also observed significant overlap in gene expression and network structure between the amniotic epithelium and published datasets of epithelial samples from the lower airway and other epithelial tissues including skin and oesophagus, suggesting a global epithelial signature.

**Conclusions:** Overall, we observed significant overlap in gene expression and network structure between paired amniotic and nasal epithelial samples, supporting the potential of the amnion as a non-invasive and abundant tissue surrogate. Observed differences between tissues were related to each tissue’s specialist functions, which remains an important consideration when assessing their overlap. Future studies aimed at investigating amnion-based biomarkers for respiratory exposures *in utero* and disease outcomes in childhood are needed to extend these results towards clinical translation.

## BACKGROUND

The amniotic epithelium offers a promising complementary tissue source for examining epithelial biology in the newborn. The amnion is a foetal-derived tissue, consisting of a single layer of cells that are typically referred to as either human amnion epithelial cells (hAECs) or human amniotic epithelial stems cells (hAESCs) [1]. These cells are pluripotent stem cells that can be differentiated into all three germ layers [1–3]. This includes the ability to differentiate into mature epithelial cells in culture, including respiratory-like epithelial cells capable of seeding lung tissue in animal models [1–3]. In addition, both the amniotic epithelium and the developing epithelium of the foetus and are exposed to the same *in utero* environment including the immune/inflammatory mediators present in amniotic fluid [4–6].

While mature epithelial tissues have been increasingly recognised for their integral function in health and disease, the amniotic epithelium has not been sufficiently studied in this context. In particular, the respiratory epithelium has been consistently identified as a key player in human respiratory disease pathogenesis, including in conditions such as allergic rhinitis [7], preterm lung disease [8], asthma [9, 10], and chronic obstructive pulmonary disease [11]. However, sampling both the upper and lower airway can be technically challenging, particularly in young children [12–16].

Therefore, we hypothesised that the amnion, when sampled from the placenta after delivery, could serve as a non-invasive and abundant source of epithelial cells and offer a viable alternative for studying respiratory epithelial biology, while overcoming the limitations of airway sampling. We aimed to assess similarities and differences in gene expression and co-expression network structure between amniotic epithelium isolated from the placenta at birth and paired newborn nasal epithelial samples.

## METHODS

### Study Setting

The Airway Epithelium Respiratory Illnesses and Allergy (AERIAL) birth cohort study [17] is nested within the ORIGINS Project [18, 19] and collects longitudinal samples and data from recruited participants from birth to 5years.

### Sample Collection and Processing

This pilot study utilised 85 matched amniotic membrane biopsies and newborn nasal brushings from the AERIAL cohort.

Matched placentae were processed within 48 hours post-birth with the amniotic membrane manually separated from the chorion and sectioned into strips [17, 20]. Nasal epithelial cells from newborns were collected within six weeks post-birth, as previously described [17]. All samples were cryopreserved in DNA/RNA Shield™ (Zymo Research) at –80°C until batch RNA extraction.

Amniotic samples were thawed, transferred into a Precellys® CK14 soft tissue ceramic bead homogenising tube (Bertin Instruments, Montigny-le-Bretonneux, France) and homogenised at 12,833xg for 30 seconds using the Precellys® 24 Tissue Homogeniser (Bertin Instruments). The homogenate was mixed with QIAzol® (1:4 v/v) and 20% (v/v) chloroform, centrifuged, the aqueous phase collected, and an equal volume of 70% (v/v) ethanol was added for total RNA extraction using the PureLink™ mini RNA extraction kit (Thermo Fisher Scientific, Waltham, MA, USA) as per manufacturer’s instructions.

Total RNA from newborn nasal brushing was extracted using Zymo Quick-RNA™ Microprep Kit (Irvine, CA, USA Research). Nasal samples were thawed, treated with proteinase K for 1hour at room temperature, and centrifuged. The supernatant was mixed with RNA Lysis buffer (1:3 v/v) and equal volume of 100% (v/v) ethanol. Samples were spun through the columns and buffers added as per manufacturer’s instructions with the following alterations: An extra 400 µL Wash Buffer spin was performed before the addition of DNase I. The DNase I provided in the kit was replaced with PureLink™ DNase I Reaction Mix and 80 µL was to samples incubated for 30 minutes. Elution was in 15 µL of DNase/RNase-free water.

For all samples, total RNA purity was determined via NanoDrop™ 1000 spectrophotometer (A260/280 > 2.0), yield by Qubit fluorometer and quality determined using the Agilent RNA 6000 Nano Kit and Agilent 2100 Bioanalyser, according to manufacturer’s instructions (Agilent Technologies, Santa Clara, CA, USA).

Extracted amnion and nasal RNA samples were sent to the Australian Genome Research Facility (AGRF) for whole-transcriptome library preparation and sequencing, using an Illumina NovaSeq 6000 (Illumina, San Diego, CA) platform generating pair-end reads (150 bp).

### Data analysis

The raw FASTQ files for gene expression were processed using the nf-core/rnaseq v3.12.0 pipeline on Setonix (Pawsey Supercomputer), supported by The Australian BioCommons Leadership Share (ABLeS). The pipeline was executed using Nextflow v23.04.2 [21], using the human Genome Reference Consortium Human Build 38 (GRCh38), alignment with STAR [22], and quantification with RSEM [23]. This resulted in an average of 25.9 million mapped reads per sample (range 10.7-51.6 million).

Outliers were detected independently for each tissue based on the standardised connectivity (Z.K) approach from the WGCNA authors with a outlier threshold of –2 [24]. In total, five nasal sample and two amnion samples were identified as outliers using this method and were removed leaving 163 samples (156-paired) for downstream analysis.

“Expressed” genes in each tissue were defined using a count per million threshold corresponding to an absolute count of at least 10 in more than 90% of samples, as previously published [14]. Asthma [25] and aberrant wound repair signatures [9] were extracted from the literature and compared to genes expressed in each tissue. Functional annotation of relevant gene lists was performed using gprofiler2, filtering out terms with more than 2000 genes [26].

Raw counts for the 11,867 genes that were defined as expressed in both tissues were transformed using the variance stabilised transformation from DESeq2 which adjusts for library size and transforms into homoscedastic data [27]. This transformed data was used as input into WGCNA co-expression network analysis [28, 29], for each tissue independently and then in combination for consensus network analysis. For all networks, a ‘signed’ network with settings deepSplit 2, mergeCutHeight 0.4, and minModuleSize 50 was used [28, 29]. A soft threshold power of 22 and 16 was chosen for amnion and nasal networks, respectively, to achieve scale free topology. Module preservation was calculated using the modulePreservation function in WGCNA using the summary preservation z-score statistic [30]. Modules with a z-score > 10 are considered strongly preserved, 2-10 are considered weakly-moderately preserved, and <2 not preserved.

To extend the data to other epithelial tissues both in an outside the airway, we extracted trancriptomic data from published datasets [14, 31]. Transcriptomic data for nasal and tracheal epithelial samples from older children were extracted from GSE118761 [14]. The Genotype-Tissue Expression (GTEx) Project was supported by the Common Fund of the Office of the Director of the National Institutes of Health, and by NCI, NHGRI, NHLBI, NIDA, NIMH, and NINDS. The data used for the analyses described in this manuscript were obtained from GTEx Analysis V8 release from the GTEx Portal on 20/05/24 [31]. Both published datasets were processed and analysed using the same methods as for this AERIAL dataset.

## RESULTS

### Comparison of gene expression profiles between amniotic and nasal epithelium

Demographic data for the included AERIAL participants are presented in Table 1. In total, 163 samples remained in the analysis after removal of outliers, with variability observed in RNA quantity (range 24–18,480ng), quality (RIN range 1.6–8.9), and mapped counts per sample (range 10.7–51.6 million) (Table 1).

**Table 1.** Demographic characteristics of the Airway Epithelium Respiratory Illnesses and Allergy (AERIAL) study.

|  |  |
| --- | --- |
| **Number of Participants** | **85** |
| **Sex** | **Female: 41**<br>**Male: 44** |
| **Maternal Ethnicity** | **Caucasian: 70**<br>**Non-Caucasian: 12**<br>**Not disclosed: 3** |
| **Method of Delivery** | **Vaginal: 41**<br>**Caesarean Section: 41**<br>**Unknown: 3** |
| **Age at Newborn Nasal Brushing (days; median, range)** | **14 (range 0 – 42)** |
| **RNA Quantity (ng)** | **Amnion: 3938 (range 119–18,480)**<br>**Nasal: 480 (range 24–2,148)** |
| **RNA Integrity Number** | **Amnion: 4.7 (range 1.6 – 8.8)**<br>**Nasal: 6.8 (range 4.8 – 8.9)** |
| **RNA-seq counts per sample (million)** | **Amnion: 23.7 (range 10.7 – 44.4)**<br>**Nasal: 28.2 (range 12.7 – 51.6)** |

To characterise the relationship between amniotic and nasal epithelial samples, we first performed principal component analysis, which as expected revealed tissue as the primary source of variation in the data, with the first principal component accounting for 80% of the variation in the dataset (Figure 1A). We next compared the overlap of expressed genes between the two tissues (see Methods). We identified 12,325 and 14,430 genes expressed in amniotic and nasal epithelium, respectively, of which 11,867 (79.7%) genes were expressed in both sites (Figure 1B). Pathway overrepresentation analysis on the 2,563 genes uniquely expressed in nasal epithelium identified terms related to ciliated cells (e.g., “axoneme assembly”), while the 458 genes uniquely expressed in amniotic epithelium were enriched for terms related to other developmental processes (e.g., “circulatory system development” and “embryonic morphogenesis”), consistent with the known specialist roles of these tissues. Overall, gene expression was modestly correlated between amniotic and nasal epithelium (Kendall’s tau = 0.54; p-value <2.2×10^−16^) (Figure 1C).

**Figure 1.**
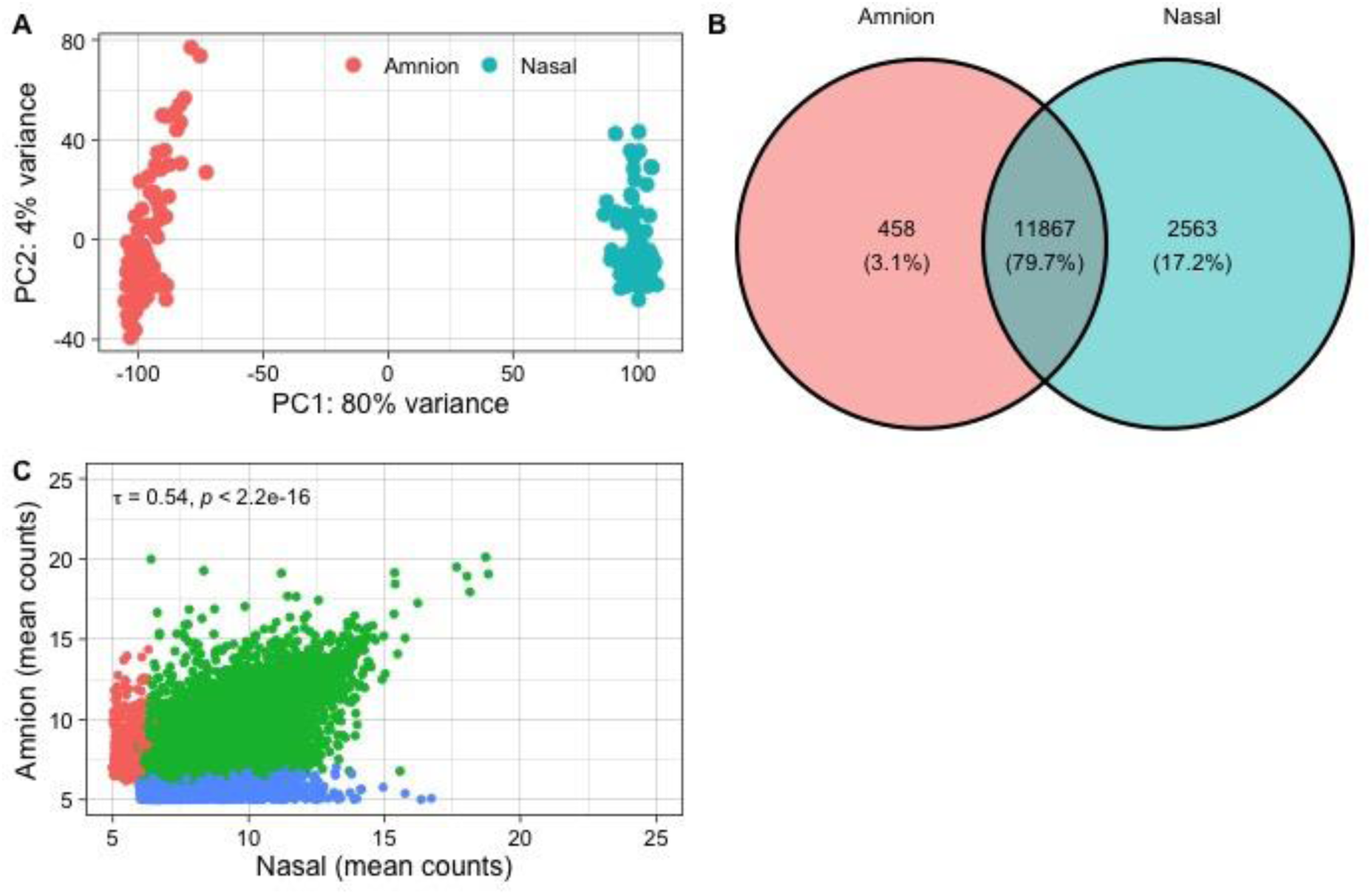
Conservation of global gene expression between amniotic (red) and nasal (blue) epithelium. (A) Principal component analysis of DESeq2 variance stabilised transformation count data. (B) Venn Diagram showing overlap in expressed genes in each tissue. Genes were defined as “expressed” in each tissue separately using a count per million threshold corresponding to an absolute count of at least 10 in more than 90% of each tissues samples. (C) Correlation plot of the mean variance stabilised transformation counts in each tissue.

We then assessed the relative expression patterns of previously published epithelial markers of asthma [25] and abnormal epithelial wound repair [9] between the two tissues. Amongst the 91 gene asthma signature, 65 genes (71.4%, hypergeometric p-value 0.022) were expressed in both amnion and nasal epithelium, while 16 genes were expressed only in nasal epithelium, 3 in amniotic epithelium, and 7 in neither tissue. Amongst the 1060 abnormal wound repair genes, 771 (72.7%, hypergeometric p-value 5.4×10^−8^) were expressed in both amnion and nasal epithelium, while 167 genes were expressed only in nasal epithelium, 38 only in amniotic epithelium, and 84 in neither tissue.

### Conservation of gene co-expression networks between amniotic and nasal epithelium

To identify preserved patterns of gene expression between amniotic and nasal epithelial samples, we performed weighted gene co-expression analysis (WGCNA) on each tissue in parallel using the 11,867 genes expressed in both tissues as input. We identified 10 network modules within the amniotic epithelium samples (Table 2), and 11 modules within the nasal epithelium samples (Table 3). Overall, we observed strong overlap in network structure with high summary preservation z-scores, with most amnion networks corresponding to one or more nasal networks and vice versa (Figure 2, Tables 2, 3). This included two network modules related to the immune response in both tissues (Tables 2,3). The exceptions were the “pink” network module from the amniotic epithelium (summary preservation z-scores 5.6) and the “turquoise” network modules from the nasal epithelium (z-score 4.2), which showed only moderate preservation. The “pink” amnion network was enriched with genes involved in the hypoxia response (e.g., “cellular response to hypoxia”; Table 2), while the “turquoise” nasal network was enriched for genes related to cilia (e.g., “cilium organisation”; Table 3).

**Figure 2.**
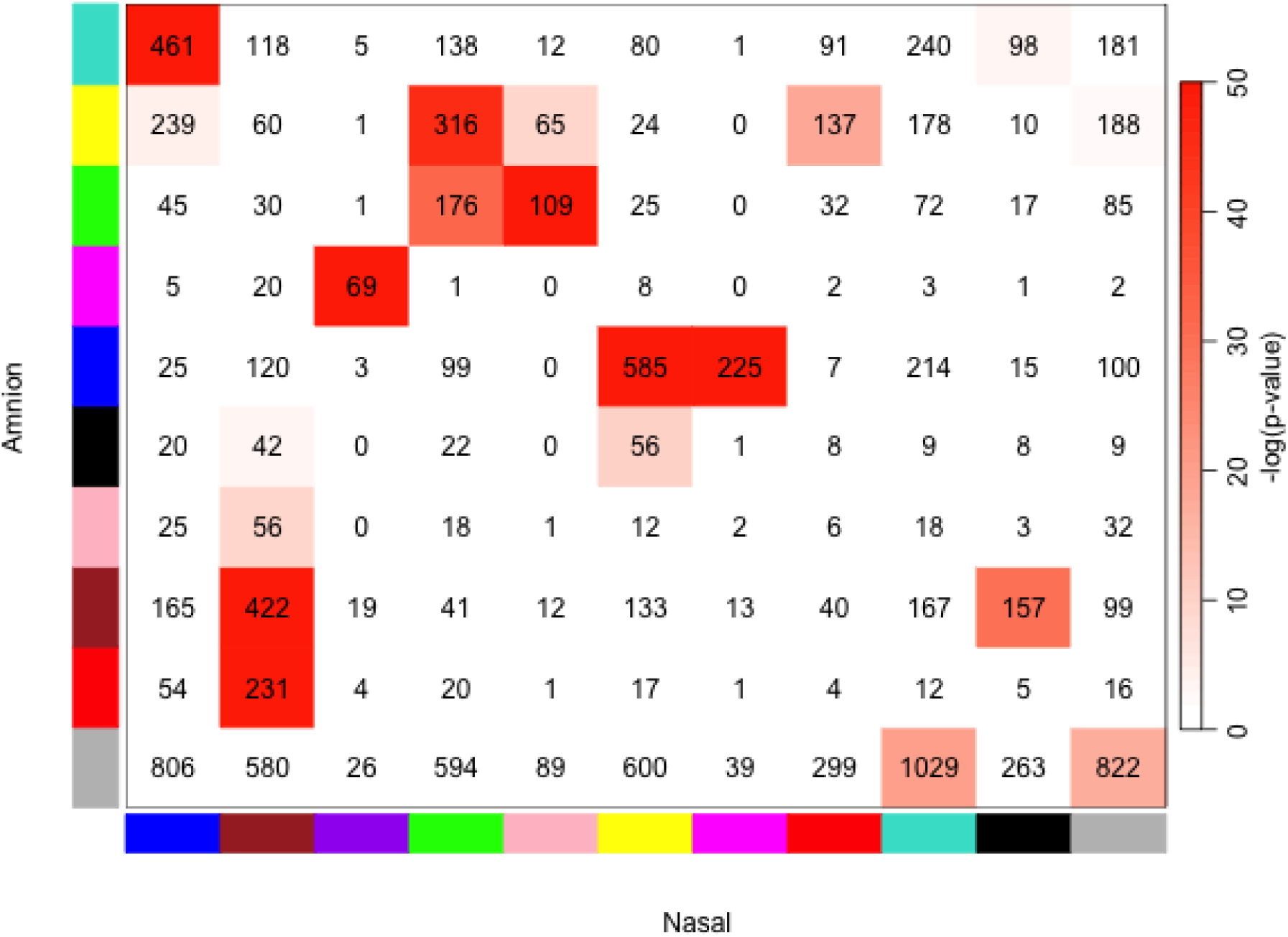
Overlap in network gene membership between amnion (y-axis) and nasal (x-axis) weighted co-expression network modules. Numeric values presented are the number of genes in common, while the red colour denotes the –log(p-value) from a Fisher’s exact test.

**Table 2.** Weighted gene co-expression network modules identified in amnion samples. The top 5 generalised biological functions for genes in each network module were identified using gprofiler2. The preservation summary z-score was calculated for the identified amnion network modules in the paired nasal samples.

| Amnion Network Module |  | Generalised Biological Function | Module Size | Preservation Z-score Summary in Nasal |
| --- | --- | --- | --- | --- |
| 1 | black | positive regulation of RNA metabolic process, protein-DNA complex organization, positive regulation of DNA-templated transcription, positive regulation of RNA biosynthetic process, chromatin organization | 175 | 12.2 |
| 2 | blue | organelle assembly, cilium organization, cilium assembly, protein-DNA complex organization, chromatin organization | 1393 | 47.1 |
| 3 | brown | cell migration, cell motility, locomotion, positive regulation of multicellular organismal process, intracellular signaling cassette | 1268 | 17.8 |
| 4 | green | cytoplasmic translation, translation, peptide biosynthetic process, peptide metabolic process, amide biosynthetic process | 592 | 21.9 |
| 5 | magenta | defense response to other organism, defense response to symbiont, response to other organism, response to external biotic stimulus, innate immune response | 111 | 40.6 |
| 6 | pink | cellular response to hypoxia, response to hypoxia, cellular response to decreased oxygen levels, response to decreased oxygen levels, cellular response to oxygen levels | 173 | 5.6 |
| 7 | red | defence response, inflammatory response, response to cytokine, cellular response to cytokine stimulus, response to biotic stimulus | 365 | 38.9 |
| 8 | turquoise | cytoskeleton organization, regulation of cellular component biogenesis, actin filament-based process, epidermis development, actin cytoskeleton organization | 1425 | 15.6 |
| 9 | yellow | aerobic respiration, oxidative phosphorylation, cellular respiration, proton motive force-driven ATP synthesis, generation of precursor metabolites and energy | 1218 | 16.9 |
| 10 | grey | *Contains genes not belonging to a specific module | 5147 | - |

**Table 3.** Weighted gene co-expression network modules identified in nasal samples. The top 5 generalised biological functions for genes in each network module were identified using gprofiler2. The preservation summary z-score was calculated for the identified nasal network modules in the paired amnion samples.

| Nasal network Module |  | Generalised Biological Function | Module Size | Summary Preservation Z-score in Amnion |
| --- | --- | --- | --- | --- |
| 1 | black | cell adhesion, enzyme-linked receptor protein signaling pathway, neurogenesis, generation of neurons, cell migration | 577 | 15.6 |
| 2 | blue | intracellular transport, establishment of localization in cell, vesicle-mediated transport, protein transport, establishment of protein localization | 1845 | 21.7 |
| 3 | brown | regulation of immune system process, intracellular signaling cassette, regulation of intracellular signal transduction, cell activation, positive regulation of immune system process | 1679 | 27.7 |
| 4 | green | translation, peptide biosynthetic process, peptide metabolic process, amide biosynthetic process, ribosome biogenesis | 1425 | 31.3 |
| 5 | magenta | regulation of mRNA metabolic process, regulation of RNA splicing, regulation of mRNA processing | 282 | 40.3 |
| 6 | pink | cytoplasmic translation, translation, peptide biosynthetic process, peptide metabolic process, amide biosynthetic process | 289 | 25.9 |
| 7 | purple | defense response to other organism, defense response to symbiont, innate immune response, response to other organism, response to external biotic stimulus | 128 | 36.5 |
| 8 | red | organonitrogen compound biosynthetic process, carbohydrate derivative biosynthetic process, mitochondrial transport, cellular lipid metabolic process, lipid metabolic process | 626 | 20.9 |
| 9 | turquoise | cilium organization, cilium assembly, plasma membrane bounded cell projection assembly, microtubule-based process, cell projection assembly | 1942 | 4.2 |
| 10 | yellow | DNA damage response, DNA metabolic process, DNA repair, cellular catabolic process, interstrand cross-link repair | 1540 | 34.2 |
| 11 | grey | *Contains genes not belonging to a specific module | 1534 | - |

Given the similarities between the amniotic and nasal network modules, we next built a consensus network to identify robust gene co-expression patterns common to both epithelial tissues. The consensus network contained ten modules of co-expressed genes (Figure 3A). Of these, six were associated with generalised cellular processes (“black”, “blue”, “brown”, “green”, “pink”, and “red”), while two were associated with immune responses (“magenta” and “turquoise”) and one with epithelial development (“yellow”)(Table 4). To assess the relative expression patterns of module genes between tissues, the median log2 fold-change difference between amnion and nasal epithelium was calculated across all genes in each module (Figure 3B). Only the “black” consensus network module (“smoothened signaling pathway”) had an absolute log2 fold-change greater than 1 (fold change –1.07), supporting an overall similar pattern of gene expression between tissues within the conserved modules.

**Figure 3.**
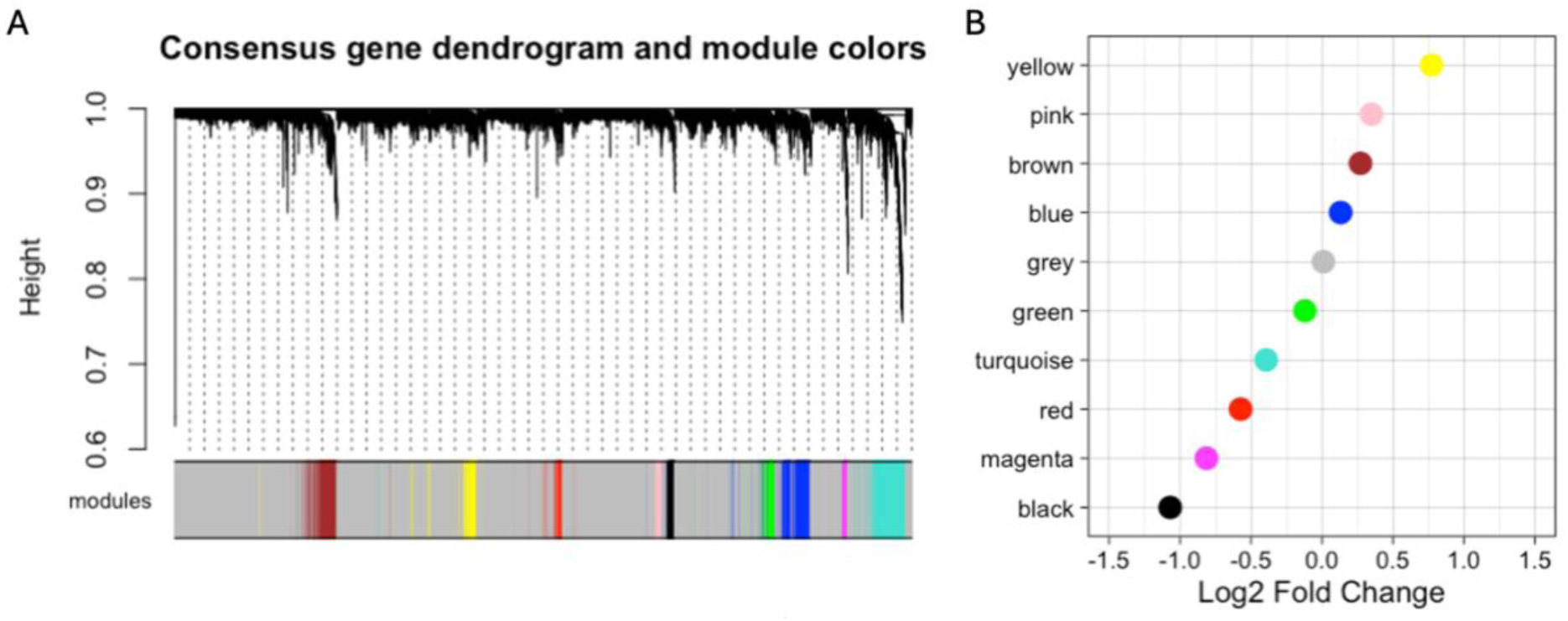
(A) Consensus weighted gene co-expression network dendrogram from amnion and nasal samples. (B) Median Log2 fold change in network module gene expression between the nasal and amnion samples.

**Table 4.** Consensus weighted gene co-expression network modules identified in paired amnion and nasal samples. The top 5 generalised biological functions for genes in each network module were identified using gprofiler2. Differential gene expression analysis was performed between amnion and nasal samples, with the median log2 fold change presented across the genes in each network module.

| Consensus Network Modules |  | Generalised Biological Function | Module Size | Median Log2 Fold Change |
| --- | --- | --- | --- | --- |
| 1 | black | smoothened signaling pathway | 118 | -1.07 |
| 2 | blue | organelle assembly, cilium organization, cilium assembly, microtubule-based process, cell cycle process | 473 | 0.132 |
| 3 | brown | cytoplasmic translation, translation, peptide biosynthetic process, organonitrogen compound biosynthetic process, peptide metabolic process | 427 | 0.272 |
| 4 | green | cilium organization, cilium assembly, plasma membrane bounded cell projection assembly, cell projection assembly, organelle assembly | 182 | -0.120 |
| 5 | magenta | defense response to virus, response to virus, response to other organism, response to external biotic stimulus, defense response to symbiont | 76 | -0.813 |
| 6 | pink | extracellular matrix organization, extracellular structure organization, external encapsulating structure organization, cell migration, tube development | 112 | 0.349 |
| 7 | red | microtubule-based movement, microtubule-based process, microtubule-based transport, cilium organization, cilium assembly | 120 | -0.573 |
| 8 | turquoise | immune response, regulation of immune system process, defense response, cell activation, leukocyte activation | 596 | -0.393 |
| 9 | yellow | epidermis development, skin development, epidermal cell differentiation, keratinocyte differentiation, epithelial cell differentiation | 241 | 0.773 |
| 10 | grey | *Contains genes not belonging to a specific module | 9522 | 0.0102 |

### Global epithelial gene signatures in the amnion

Given that the amniotic epithelium is comprised of stem cells capable of differentiating into all mature epithelial tissues in the body, we next queried the expression of global epithelial markers using cell type enrichment analysis with annotations from the human protein atlas. While many of the consensus co-expression network modules were enriched for genes annotated to the nasopharynx (i.e. “blue”, “brown”, “red”, and “yellow” consensus modules) as expected, we also observed significant enrichment in consensus module genes that were annotated to the lower airways (e.g., lung/bronchus) as well as other epithelial tissues including skin, gastrointestinal tract (e.g., oesophagus/intestine) and genitourinary tract (e.g., kidney/vagina).

To explore consistency with the unified airway hypothesis [14], we next assessed for conservation in gene expression and co-expression network structure with our previously published epithelial dataset of matched nasal and tracheal epithelium in older children [14]. Consistent with this hypothesis [14], 11,392 of the 11,867 genes (96.0%) expressed in both tissues in the current study were also expressed in both nasal and tracheal epithelium in an older children dataset. Overall, the consensus network structure was also strongly preserved within nasal and tracheal samples (Table 5). We also performed the same analyses in the epithelial-dominant samples from the adult GTEx dataset [31], and identified a high-degree of overlap in gene expression between our study and the published lung (11,124/11,867; 93.7%), skin (11,124/11,867; 93.7%), and oesophagus (11,107/11,867; 93.6%) samples, with overall strong preservation of network structure (Table 5).

**Table 5.** Preservation of consensus weighted gene co-expression network modules from our study within other epithelial datasets from the published literature. Modules with no (z-score <2) or moderate preservation (z-score 2-10) are flagged with an *.

|  |  | Summary Preservation Z-Score |  |  |  |  |
| --- | --- | --- | --- | --- | --- | --- |
| Consensus Network Modules |  | GSE118761 Nasal | GSE118761 Tracheal | GTEX Lung | GTEX Skin | GTEX Oesophagus |
| 1 | black | 19.3 | 11.9 | 4.6* | 9.3* |  |
| 2 | blue | 22.4 | 18.7 | 12.5 | 27.3 |  |
| 3 | brown | 25.0 | 32.5 | 21.0 | 51.3 |  |
| 4 | green | 39.6 | 33.6 | 13.8 | 10.4 |  |
| 5 | magenta | 42.7 | 40.1 | 10.3 | 14.9 |  |
| 6 | pink | 5.5* | 5.8* | 8.1* | 18.1 |  |
| 7 | red | 38.2 | 32.9 | 10.5 | 1.4* |  |
| 8 | turquoise | 39.1 | 54.3 | 12.7 | 19.0 |  |
| 9 | yellow | 18.2 | 8.6* | 6.8* | 18.6 |  |
| 10 | grey | - | - | - | - | - |

## DISCUSSION

The airway epithelium plays a central role in the pathogenesis of respiratory disease, but investigations into epithelial physiology are limited by the challenges of sampling airway tissue. We hypothesised that the amniotic epithelium may serve as a useful tissue surrogate for characterising aspects of the respiratory epithelium in the newborn. In support of this hypothesis, we observed significant overlap in global gene expression, expression of epithelial biomarkers of asthma/wound repair, and gene co-expression network structure between paired amniotic and nasal epithelial samples.

To our knowledge, this is the first study to compare gene expression patterns in the amniotic and respiratory epithelium. However, previous studies have highlighted the relevance of the amnion to understand the effect of pregnancy conditions, such as pre-eclampsia, on the foetus [5, 32, 33]. In general, the prenatal environment has been increasingly recognised as a critical factor in the origins of chronic respiratory disease in children and adults such as asthma and chronic obstructive pulmonary disease (COPD) [34, 35]. For example, prenatal exposures (e.g., cigarette smoke, pollution, diet) and poorly controlled maternal asthma during pregnancy significantly increased risk of asthma development and respiratory morbidity throughout life [4, 34, 36, 37]. Given that the amniotic and respiratory epithelium of the foetus are both in direct contact with amniotic fluid, which contains immune mediators such as cytokines and chemokines [6], it is likely the epithelial cells in both tissues would undergo similar transcriptional changes to these factors. Unfortunately, critical exposures such as cigarette smoking during pregnancy were rare in this study (n=4), and so we did not have the power to quantify their shared influence. While our study supports global conservation of many core biological processes between tissues, additional work is required to test whether biomarkers of *in utero* exposures are similarly conserved between tissues.

One of the key roles of the respiratory epithelium that mechanistically link it to respiratory disease pathogenesis is its role in directing airway immune responses to inhaled pathogens [38]. In keeping with this function, we observed multiple co-expression network modules in the nasal samples that were enriched for genes related to innate immunity and antimicrobial defence responses, as expected. Reassuringly, these network modules exhibited strong preservation statistics in the amnion samples and these terms were also enriched in the “turquoise” and “magenta” network modules from the consensus co-expression network analysis. Another central epithelial process that has been linked to respiratory disease development by our group and others is defective wound repair [9, 39–46]. This epithelial endotype is characterised by aberrant cell migration and wound repair due to extracellular matrix dysregulation [42, 47], abnormal barrier function [45], and impaired innate immunity to viral infection [41, 43], and associated with higher respiratory morbidity [9, 39, 40, 44]. Importantly, 72.7% of genes significantly differentially expressed in basal epithelial cells from children with and without abnormal wound repair [9], were also expressed in both nasal and amniotic epithelial cells in this study. Similarly, genes associated with asthma in a nasal epithelial transcriptomic meta-analysis [25] were also largely expressed in both nose and amnion. Collectively, these data suggest the amnion may serve as a useful surrogate for assessing core airway functions of the respiratory epithelium.

While a large degree of overlap in gene expression and network structure was observed between tissues, significant differences were still observed as expected given the different functions of amniotic and nasal epithelium. In keeping with the stem cell characteristics of the amnion [1, 3], genes uniquely expressed in the amniotic epithelium were enriched for terms related to morphogenesis and development. Similarly, genes and network modules uniquely identified in the nose were enriched for genes and processes related to ciliated cells, reflecting the terminal differentiation of the nasal epithelium into specialised airway cells (e.g. multiciliated epithelial cells) [13]. While the overlap in biologically relevant processes related to immunity, wound repair, and asthma support the potential use of the amnion as a surrogate, these differences between tissues in terms of cellular composition, function, and gene expression will remain an important consideration when studying epithelial biology or developing predictive biomarkers in each tissue.

While this study was focused on assessing the ability of the amniotic epithelium to serve as a surrogate for the nasal epithelium, the ability of the amniotic epithelium stem-like cells to differentiate to all mature epithelium in the body, would supports broader surrogacy of the amnion for other epithelial tissues [1, 3]. In keeping with this, we observed preservation of network structure and gene expression between nose/amnion in this study with other previously published epithelial tissue datasets including lung, skin, and oesophagus [14, 31]. Similarly, many of the conserved network modules identified in our dataset were enriched for human protein atlas terms related to other epithelial tissues, suggesting a strong global epithelial signature present in the amnion. Given that many epithelial tissues including those in the genitourinary and gastrointestinal tract have similar challenges to the respiratory epithelium in terms of sampling difficulty, the amnion may additionally serve as a non-invasive tissue surrogate for investigating global epithelial biology at birth.

Our study has some limitations to consider. In this study we performed bulk tissue RNA-seq to investigate the overlap in global gene expression in each tissue. However, given the known cellular composition differences between tissues, future studies performing single cell sequencing would provide additional information about the complexity of each tissue and overlap in gene expression patterns [31, 48]. In addition, due to the introduction of COVID-19 pandemic restrictions which limited hospital access for research [17], the AERIAL study was changed so that newborn nasal sampling was performed at home and up to six weeks of age. Given that gene expression in the nasal epithelium will be modulated by environmental exposures postnatally, this enforced change likely introduced a source of variability into the dataset that needs to be considered. Finally, given the inherent challenges of sampling the newborn nasal epithelium, we observed a wide range of RNA quality and quantity that may have partially influence the results of this study. However, even with these additional sources of variation, the observation of a relevant overlap between tissues independent of composition, sampling time, and RNA quality/quantity supports the potential for surrogacy.

In summary, we have demonstrated significant overlap in gene expression and co-expression network structure to support the amniotic epithelium as a meaningful surrogate for a newborn’s respiratory epithelium. Future studies aimed at identifying biomarkers of *in utero* exposures and/or future respiratory disease risk are needed to extend these observations through to clinical translation.

## DECLARATIONS

### Ethics approval and consent to participate

This study was performed in accordance with the Declaration of Helsinki. This human study was approved by Ramsay Health Care WA-SA HREC – approval: #1908. All parents, guardians, or next of kin provided written informed consent for the newborn participants to participate in this study.

### Consent for publication

Not applicable

### Availability of data and materials

The datasets generated and/or analysed during the current study are available in the Gene Expression Omnibus repository.

### Authors’ contributions

DGH, PAR, TI, YVK, GZ, AK, and SMS developed the methods and analysis plan. DJM, AK, and SMS designed the original AERIAL cohort. EK-S managed the AERIAL cohort. EK-S, JL, and TI collected the data. DGH and PAR performed the data analysis. DGH drafted the manuscript. PAR, EK-S, JL, TI, YVK, GZ, DJM, AK, and SMS reviewed and revised the manuscript. All authors read and approved the final manuscript.

## Acknowledgements.

We would like to thank the contribution of the AERIAL families, together with the cohort study investigators not listed as authors: Jose A Caparros-Martin, Peter Le Souef, Anthony Bosco, Susan Prescott, Desiree Silva and Asha Bowen. We also extend our gratitude to the hardworking and the dedicated research team, for the recruitment, liaising and sample collection over the duration of the cohort study, including the AERIAL Laboratory team: Minda Amin, Bailee Renouf, Courtney Kidd, and Ashleigh Heng-Chin.

AERIAL is a sub-project of The ORIGINS Project. This unique long-term study, a collaboration between Telethon Kids Institute and Joondalup Health Campus, is one of the most comprehensive studies of pregnant women and their families in Australia to date, recruiting 10,000 families over a decade from the Joondalup and Wanneroo communities of Western Australia. We are grateful to all the ORIGINS families who support the project. We would also like to acknowledge and thank the following teams and individuals who have made The ORIGINS Project possible: The ORIGINS Project team; Joondalup Health Campus (JHC); members of ORIGINS Community Reference and Participant Reference Groups; Research Interest Groups and the ORIGINS Scientific Committee; Telethon Kids Institute; City of Wanneroo; City of Joondalup; and Professor Fiona Stanley. The ORIGINS Project has received core funding support from the Telethon Perth Children’s Hospital Research Fund, Joondalup Health Campus, the Paul Ramsay Foundation, and the Commonwealth Government of Australia through the Channel 7 Telethon Trust. Substantial in-kind support has been provided by Telethon Kids Institute and Joondalup Health Campus.

## REFERENCES

1. Qiu C, Ge Z, Cui W, Yu L, Li J. Human Amniotic Epithelial Stem Cells: A Promising Seed Cell for Clinical Applications. Int J Mol Sci 2020, 21(20).

2. Moodley Y, Ilancheran S, Samuel C, Vaghjiani V, Atienza D, Williams ED, Jenkin G, Wallace E, Trounson A, Manuelpillai U. Human amnion epithelial cell transplantation abrogates lung fibrosis and augments repair. Am J Respir Crit Care Med 2010, 182(5):643–651.

3. Pratama G, Vaghjiani V, Tee JY, Liu YH, Chan J, Tan C, Murthi P, Gargett C, Manuelpillai U. Changes in culture expanded human amniotic epithelial cells: implications for potential therapeutic applications. PLoS One 2011, 6(11):e26136.

4. Liu X, Agerbo E, Schlunssen V, Wright RJ, Li J, Munk-Olsen T. Maternal asthma severity and control during pregnancy and risk of offspring asthma. J Allergy Clin Immunol 2018, 141(3):886–892 e883.

5. Suzuki M, Maekawa R, Patterson NE, Reynolds DM, Calder BR, Reznik SE, Heo HJ, Einstein FH, Greally JM. Amnion as a surrogate tissue reporter of the effects of maternal preeclampsia on the fetus. Clin Epigenetics 2016, 8:67.

6. Weissenbacher T, Laubender RP, Witkin SS, Gingelmaier A, Schiessl B, Kainer F, Friese K, Jeschke U, Dian D, Karl K. Influence of maternal age, gestational age and fetal gender on expression of immune mediators in amniotic fluid. BMC Res Notes 2012, 5:375.

7. Berni Canani R, Caminati M, Carucci L, Eguiluz-Gracia I. Skin, gut, and lung barrier: Physiological interface and target of intervention for preventing and treating allergic diseases. Allergy 2024, 79(6):1485–1500.

8. Looi K, Evans DJ, Garratt LW, Ang S, Hillas JK, Kicic A, Simpson SJ. Preterm birth: Born too soon for the developing airway epithelium? Paediatr Respir Rev 2019, 31:82–88.

9. Iosifidis T, Sutanto EN, Buckley AG, Coleman L, Gill EE, Lee AH, Ling KM, Hillas J, Looi K, Garratt LW et al. Aberrant cell migration contributes to defective airway epithelial repair in childhood wheeze. JCI Insight 2020, 5(7).

10. Iosifidis T, Sutanto EN, Montgomery ST, Agudelo-Romero P, Looi K, Ling KM, Shaw NC, Garratt LW, Hillas J, Martinovich KM et al. Dysregulated Notch Signaling in the Airway Epithelium of Children with Wheeze. J Pers Med 2021, 11(12).

11. Love ME, Proud D. Respiratory Viral and Bacterial Exacerbations of COPD-The Role of the Airway Epithelium. Cells 2022, 11(9).

12. Baschal EE, Larson ED, Bootpetch Roberts TC, Pathak S, Frank G, Handley E, Dinwiddie J, Moloney M, Yoon PJ, Gubbels SP et al. Identification of Novel Genes and Biological Pathways That Overlap in Infectious and Nonallergic Diseases of the Upper and Lower Airways Using Network Analyses. Front Genet 2019, 10:1352.

13. Deprez M, Zaragosi LE, Truchi M, Becavin C, Ruiz Garcia S, Arguel MJ, Plaisant M, Magnone V, Lebrigand K, Abelanet S et al. A Single-Cell Atlas of the Human Healthy Airways. Am J Respir Crit Care Med 2020, 202(12):1636–1645.

14. Kicic A, de Jong E, Ling KM, Nichol K, Anderson D, Wark PAB, Knight DA, Bosco A, Stick SM, Waerp et al. Assessing the unified airway hypothesis in children via transcriptional profiling of the airway epithelium. J Allergy Clin Immunol 2020, 145(6):1562–1573.

15. Miller D, Turner SW, Spiteri-Cornish D, McInnes N, Scaife A, Danielian PJ, Devereux G, Walsh GM. Culture of airway epithelial cells from neonates sampled within 48-hours of birth. PLoS One 2013, 8(11):e78321.

16. Stokes AB, Kieninger E, Schogler A, Kopf BS, Casaulta C, Geiser T, Regamey N, Alves MP. Comparison of three different brushing techniques to isolate and culture primary nasal epithelial cells from human subjects. Exp Lung Res 2014, 40(7):327–332.

17. Kicic-Starcevich E, Hancock DG, Iosifidis T, Agudelo-Romero P, Caparros-Martin JA, Karpievitch YV, Silva D, Turkovic L, Le Souef PN, Bosco A et al. Airway epithelium respiratory illnesses and allergy (AERIAL) birth cohort: study protocol. Front Allergy 2024, 5:1349741.

18. D’Vaz N, Kidd C, Miller S, Amin M, Davis JA, Talati Z, Silva DT, Prescott SL. The ORIGINS Project Biobank: A Collaborative Bio Resource for Investigating the Developmental Origins of Health and Disease. Int J Environ Res Public Health 2023, 20(13).

19. Silva DT, Hagemann E, Davis JA, Gibson LY, Srinivasjois R, Palmer DJ, Colvin L, Tan J, Prescott SL. Introducing the ORIGINS project: a community-based interventional birth cohort. Rev Environ Health 2020, 35(3):281–293.

20. Magatti M, Caruso M, De Munari S, Vertua E, De D, Manuelpillai U, Parolini O. Human Amniotic Membrane-Derived Mesenchymal and Epithelial Cells Exert Different Effects on Monocyte-Derived Dendritic Cell Differentiation and Function. Cell Transplant 2015, 24(9):1733–1752.

21. Di Tommaso P, Chatzou M, Floden EW, Barja PP, Palumbo E, Notredame C. Nextflow enables reproducible computational workflows. Nat Biotechnol 2017, 35(4):316–319.

22. Dobin A, Davis CA, Schlesinger F, Drenkow J, Zaleski C, Jha S, Batut P, Chaisson M, Gingeras TR. STAR: ultrafast universal RNA-seq aligner. Bioinformatics 2013, 29(1):15–21.

23. Li B, Dewey CN. RSEM: accurate transcript quantification from RNA-Seq data with or without a reference genome. BMC Bioinformatics 2011, 12:323.

24. Oldham MC, Langfelder P, Horvath S. Network methods for describing sample relationships in genomic datasets: application to Huntington’s disease. BMC Syst Biol 2012, 6:63.

25. Sajuthi SP, Everman JL, Jackson ND, Saef B, Rios CL, Moore CM, Mak ACY, Eng C, Fairbanks-Mahnke A, Salazar S et al. Nasal airway transcriptome-wide association study of asthma reveals genetically driven mucus pathobiology. Nat Commun 2022, 13(1):1632.

26. Kolberg L, Raudvere U, Kuzmin I, Adler P, Vilo J, Peterson H. g:Profiler-interoperable web service for functional enrichment analysis and gene identifier mapping (2023 update). Nucleic Acids Res 2023, 51(W1):W207–W212.

27. Love MI, Huber W, Anders S. Moderated estimation of fold change and dispersion for RNA-seq data with DESeq2. Genome Biol 2014, 15(12):550.

28. Langfelder P, Horvath S. WGCNA: an R package for weighted correlation network analysis. BMC Bioinformatics 2008, 9:559.

29. Langfelder P, Horvath S. Fast R Functions for Robust Correlations and Hierarchical Clustering. J Stat Softw 2012, 46(11).

30. Langfelder P, Luo R, Oldham MC, Horvath S. Is my network module preserved and reproducible? PLoS Comput Biol 2011, 7(1):e1001057.

31. Consortium GT. The Genotype-Tissue Expression (GTEx) project. Nat Genet 2013, 45(6):580–585.

32. Ching T, Song MA, Tiirikainen M, Molnar J, Berry M, Towner D, Garmire LX. Genome-wide hypermethylation coupled with promoter hypomethylation in the chorioamniotic membranes of early onset pre-eclampsia. Mol Hum Reprod 2014, 20(9):885–904.

33. Kim J, Pitlick MM, Christine PJ, Schaefer AR, Saleme C, Comas B, Cosentino V, Gadow E, Murray JC. Genome-wide analysis of DNA methylation in human amnion. ScientificWorldJournal 2013, 2013:678156.

34. Altman MC, Calatroni A, Ramratnam S, Jackson DJ, Presnell S, Rosasco MG, Gergen PJ, Bacharier LB, O’Connor GT, Sandel MT et al. Endotype of allergic asthma with airway obstruction in urban children. J Allergy Clin Immunol 2021.

35. Krauss-Etschmann S, Bush A, Bellusci S, Brusselle GG, Dahlen SE, Dehmel S, Eickelberg O, Gibson G, Hylkema MN, Knaus P et al. Of flies, mice and men: a systematic approach to understanding the early life origins of chronic lung disease. Thorax 2013, 68(4):380–384.

36. Wada T, Adachi Y, Murakami S, Ito Y, Itazawa T, Tsuchida A, Matsumura K, Hamazaki K, Inadera H, Japan E et al. Maternal exposure to smoking and infant’s wheeze and asthma: Japan Environment and Children’s Study. Allergol Int 2021, 70(4):445–451.

37. Zazara DE, Wegmann M, Giannou AD, Hierweger AM, Alawi M, Thiele K, Huber S, Pincus M, Muntau AC, Solano ME et al. A prenatally disrupted airway epithelium orchestrates the fetal origin of asthma in mice. J Allergy Clin Immunol 2020.

38. Knight DA, Holgate ST. The airway epithelium: structural and functional properties in health and disease. Respirology 2003, 8(4):432–446.

39. Bizzintino J, Lee WM, Laing IA, Vang F, Pappas T, Zhang G, Martin AC, Khoo SK, Cox DW, Geelhoed GC et al. Association between human rhinovirus C and severity of acute asthma in children. Eur Respir J 2011, 37(5):1037–1042.

40. Hillas J, Evens D, Hemy N, Iosifidis T, Looi K, Garratt L, Simpson S, Kicic A. Neonatal predictors of aberrant wound repair in very preterm infants. Respirology 2019, 24:103.

41. Kicic A, Sutanto EN, Stevens PT, Knight DA, Stick SM. Intrinsic biochemical and functional differences in bronchial epithelial cells of children with asthma. Am J Respir Crit Care Med 2006, 174(10):1110–1118.

42. Kicic A, Hallstrand TS, Sutanto EN, Stevens PT, Kobor MS, Taplin C, Pare PD, Beyer RP, Stick SM, Knight DA. Decreased fibronectin production significantly contributes to dysregulated repair of asthmatic epithelium. Am J Respir Crit Care Med 2010, 181(9):889–898.

43. Kicic A, Stevens PT, Sutanto EN, Kicic-Starcevich E, Ling KM, Looi K, Martinovich KM, Garratt LW, Iosifidis T, Shaw NC et al. Impaired airway epithelial cell responses from children with asthma to rhinoviral infection. Clin Exp Allergy 2016, 46(11):1441–1455.

44. Khoo SK, Read J, Franks K, Zhang G, Bizzintino J, Coleman L, McCrae C, Oberg L, Troy NM, Prastanti F et al. Upper Airway Cell Transcriptomics Identify a Major New Immunological Phenotype with Strong Clinical Correlates in Young Children with Acute Wheezing. J Immunol 2019, 202(6):1845–1858.

45. Looi K, Buckley AG, Rigby PJ, Garratt LW, Iosifidis T, Zosky GR, Larcombe AN, Lannigan FJ, Ling KM, Martinovich KM et al. Effects of human rhinovirus on epithelial barrier integrity and function in children with asthma. Clin Exp Allergy 2018, 48(5):513–524.

46. Stevens PT, Kicic A, Sutanto EN, Knight DA, Stick SM. Dysregulated repair in asthmatic paediatric airway epithelial cells: the role of plasminogen activator inhibitor-1. Clin Exp Allergy 2008, 38(12):1901–1910.

47. Stevens PT, Kicic A, Sutanto EN, Knight DA, Stick SM. Dysregulated repair in asthmatic paediatric airway epithelial cells: the role of plasminogen activator inhibitor-1. Clinical and Experimental Allergy 2008, 38(12):1901–1910.

48. Wang WS, Lin YK, Zhang F, Lei WJ, Pan F, Zhu YN, Lu JW, Zhang CY, Zhou Q, Ying H et al. Single cell transcriptomic analysis of human amnion identifies cell-specific signatures associated with membrane rupture and parturition. Cell Biosci 2022, 12(1):64.

